# Gene augmentation and read-through rescue channelopathy in an iPSC-RPE model of congenital blindness

**DOI:** 10.1101/485847

**Authors:** Pawan K. Shahi, Dalton Hermans, Divya Sinha, Simran Brar, Hannah Moulton, Sabrina Stulo, Katarzyna D. Borys, Elizabeth Capowski, De-Ann M. Pillers, David M. Gamm, Bikash. R. Pattnaik

## Abstract

**Purpose:** Mutations in the *KCNJ13* gene are known to cause Leber’s Congenital Amaurosis (LCA16), an inherited pediatric blindness. *KCNJ13* gene encodes the Kir7.1 subunit protein which acts as a tetrameric inwardly rectifying potassium ion channel in the retinal pigment epithelium to maintain ionic homeostasis thereby allowing photoreceptors to encode visual information. We sought to determine if genetic approaches might be effective in treating blindness due to mutations in *KCNJ13*.

**Methods:** We developed patient-derived hiPSC-RPE carrying an autosomal recessive nonsense mutation in the *KCNJ13* gene (c.158G>A, p.Trp53*). We performed biochemical and electrophysiology assays of Kir7.1 function. Both small molecule read-through drug and gene-therapy approaches were tested using this disease-in-a-dish approach.

**Results:** We found that the LCA16 hiPSC-RPE had normal morphology but did not express a functional Kir7.1 channel and was unable to demonstrate normal physiology. Following read-through drug treatment, the LCA16 hiPSC cells were hyperpolarized by 30 mV and Kir7.1 current was restored. Similarly, loss-of-function of Kir7.1 channel was circumvented by lentiviral gene delivery to the hiPSC-RPE cells. In either approach, Kir7.1 protein was expressing normally with restoration of membrane potential and Kir7.1 current.

**Conclusion:** Loss-of-function mutation in Kir7.1 is a cause of LCA. Using either read-through therapy or gene augmentation, we rescued Kir7.1 channel function in patient-derived iPSC-RPE cells via a precision medicine approach.

## INTRODUCTION

Mutations in at least 21 genes expressed in the outer retina, i.e., the photoreceptors and retinal pigment epithelium (RPE), cause a severe form of early childhood inherited blindness known as Leber’s Congenital Amaurosis (LCA). Within the last decade, multiple autosomal recessive pathogenic variants of the *KCNJ13* gene (OMIM: 603208, chromosomal location: 2q37.1) have been identified in patients with the LCA phenotype (LCA16 OMIM-614186, the 16^th^ gene shown to cause LCA).^1–3^ LCA16 pathogenic variants include c.158G>A (p.Trp53*), c.359T>C (p.Ile120Thr), c.458C>T (p.Thr153Ile), c.496C>T (p.Arg166*), and c.722T>C (p.Leu241Pro).^1; 2; 4^ In addition, the compound heterozygous *KCNJ13* pathogenic variants c.314 G>T (p.Ser105Ile) and c.655C>T (p.Gln219Ter) were shown to cause early-onset retinal dystrophy in an LCA patient.^3^ An autosomal dominant *KCNJ13* pathogenic variant, c.484C>T (p.Arg162Trp), causes the early-onset blindness called snowflake vitreoretinal degeneration (SVD OMIM-193230).^5^

*KCNJ13* encodes the inwardly rectifying potassium channel, Kir7.1, and it is expressed in several tissues.^6; 7^ In the retina, Kir7.1 is localized exclusively in the apical processes of RPE, where it controls retinal function and health.^8; 9^ The major contribution of the Kir7.1 channel is in supporting the glial function of the RPE cells as modeled by LaCour.^10^ Light activation of photoreceptors reduces RPE extracellular K+ concentration from 5 mM to 2 mM. Conductance of the Kir7.1 channel is increased when extracellular K+ is decreased, and vice versa.^11^ Thus, *KCNJ13* loss-of-function mutations will directly impact K+ buffering of the tight subretinal space and thereby alter photoreceptor function. A non-functional Kir7.1 channel will cause a sustained RPE cell depolarization that lead to cell death as has been reported after *Kcnj13* gene knock-out or suppression in mice.^9; 12^ The role of the Kir7.1 channel in other organs remains to be elucidated, but to-date, all mutations show a single ophthalmologic phenotype of blindness.^13; 14^

A loss-of-function in *KCNJ13*, as with other channelopathies, is a convenient gene therapy target compared to the dominant negative SVD variant that alters wild-type channel function.^15^ Technical challenges include controlling the level of expression and to successfully translate, traffick and assemble the ion channel as a functional unit in the membrane. Similarly, misincorporation of an amino acid during the translation of a non-sense codon limits the likelihood that the synthesis of full-length protein with function will be the outcome. To address these challenges, we adopted two precision medicine approaches where we used patient-derived iPSC-RPE cells to model LCA16 and explore novel fast-track approaches to mutation-specific therapies.

## MATERIALS AND METHODS

### Ethical guidance

This study was approved by the University of Wisconsin-Madison’s Institutional Review Board in accordance with federal regulations, state laws, and local and University policies.

### Differentiation of hiPSC-RPE

Fibroblasts from two subjects were reprogrammed to induced pluripotent stem cells and cultured using established methods.^16–18^ One of the subjects was a LCA16 patient with p.Trp53* autosomal recessive mutation in the *KCNJ13* gene, and the second subject (a non-symptomatic family member) was heterozygous for this mutation. Both the wild type hiPSC cells and hfRPE cells used as controls in our experiment have been previously characterized in our laboratory.^18; 19^

The hiPSC lines were differentiated to RPE using protocols described earlier.^17; 18; 20; 21^ Briefly, hiPSCs were cultured either on mouse embryonic fibroblasts (MEFs) in iPS cell media (Dulbecco’s modified Eagle’s medium (DMEM): F12 (1:1), 20% Knockout Serum, 1% minimal essential medium (MEM) non-essential amino acids, 1% GlutaMAX, β-mercaptoethanol, 20 ng/ml FGF-2), or on Matrigel with mTeSR1 media. Cells were lifted enzymatically and grown as embryoid bodies (EBs) in iPS media without FGF-2, and at day 4, changed to neural induction media (NIM; DMEM:F12; 1% N2 supplement, 1% MEM non-essential amino acids, 1% L-Glutamine, 2 µg/ml Heparin), or in mTeSR1 and gradually transitioned to NIM by day 4. There were no differences observed in RPE differentiation between these two approaches. At day 7, free-floating EBs were plated on laminin-coated culture plates to continue differentiation as adherent culture. At day 16, the 3D neural structures were removed, and media was switched to retinal differentiation media (DMEM/F12 (3:1), 2% B27 supplement (without retinoic acid), 1% Antibiotic-Antimycotic). Remaining adhered cells were allowed to continue differentiation for an additional 45 days, followed by microdissection and passaging of pigmented RPE patches to obtain purified monolayers of RPE as described earlier.^21^ MEFs, Matrigel and FGF-2 were purchased from WiCell (Madison, WI), and all other tissue culture reagents were purchased from ThermoFisher. Karyotype experiments were performed by chromosome analysis by WiCell (Madison, WI) in accordance with general principles developed by the International Stem Cell Banking Initiative.

### RT-PCR and Restriction Fragment Length Polymorphism (RFLP)

Total RNA was isolated from mature hiPSC-RPE cells from both patient and the carrier, wild type hiPSC-RPE, and hfRPE using the RNeasy kit according to manufacturer’s instructions (Qiagen). The quality and the concentration of the isolated RNA was measured using a Nanodrop (ThermoFisher) and 200 ng of RNA was used for cDNA synthesis using the Superscript III first strand cDNA synthesis kit according to manufacturer’s instructions (ThermoFisher). PCR was performed with MyTaqHS master mix (Bioline) in a final volume of 25 µl with the following conditions: 95°C for 5 min followed by 35 cycles of denaturation at 95°C for 15 s, annealing at 55°C for 30 s, and extension at 72°C for 30 s. A final extension step was done for 10 min at 72°C and amplification products were visualized by electrophoresis on a 2% agarose gel containing Midori green advanced stain (Nippon Genetics Europe). For the RFLP assay, PCR was performed as described with primers specific to the full length *KCNJ13* mRNA (Fwd 5′-GCTTCGAATTCCGACAGCAGTAATTG-3′ and Rev 5′-ATCCGGTGGATCCTTATTCTGTCAGT-3′). The PCR products were digested by NheI restriction enzyme (ThermoFisher) and visualized by electrophoresis on a 2% agarose gel containing Midori green advanced stain (Nippon Genetics Europe).

### Transmission Electron Microscopy

Monolayers of hiPSC-RPE on transwell inserts (Corning, Cat#3470) were fixed with a solution of 2.5% glutaraldehyde, 2.0% paraformaldehyde in 0.1M sodium phosphate buffer (PB), pH 7.4 for ~1 hr at room temperature (RT). Samples were rinsed five times for 5 min in 0.1M PB. The rinsed cultures were post-fixed in 1% Osmium Tetroxide (OsO4), 1% potassium ferrocyanide in PB for 1 h at RT. Following post-fixation, samples were rinsed in PB, as before, followed by 3 × 5 min rinses in distilled water to clear the phosphates. The samples were then stained en bloc in uranyl acetate for 2 h at RT and dehydrated using an ethanol series. The membrane was cut from the transwell support, placed in an aluminum weighing dish, transitioned in propylene oxide (PO) and allowed to polymerize in fresh PilyBed 812 (Polysciences Inc. Warrington, PA). Ultrathin sections were prepared from these polymerized samples and processed before capturing and documenting the images with FEI CM120 transmission electron microscope mounted with AMT BioSprint12 (Advanced Microscopy Techniques, Corp. Woburn, MA) digital camera. All electron microscope images were subjected to morphometry analysis of mitochondria number and length of apical processes. Mitochondira were counted manually using a blind-fold approach. We used a 10 um^2 area of the image and manually counted number of mitochondria. For apical processes, we used the ImageJ program to define length of each apical process. All data from at least seven different images were averaged in Excel (Microsoft Corp., Redmond, WA, USA).

### Immunocytochemistry (ICC)

Transwell inserts with monolayer of hiPSC-RPE cells from either the patient or control were fixed as follows: the transwell membrane was cut out and fixed by immersing it in 4% paraformaldehyde in phosphate-buffered saline for 10 min in the dark. The membrane with cells was washed with chilled PBS twice and blocked for 2 h in blocking solution that contained 5% goat serum and 0.25% Tween-20 in 1X PBS. For confocal microscopy, the cells were incubated for 24-48 h with primary antibodies raised against Kir7.1 (mouse monoclonal IgG, 1:250-Santa Cruz), and ZO-1 (rabbit polyclonal, 2.5 µg/ml – ThermoFisher) prepared in incubation solution (Blocking solution diluted in 1:3 with 1X PBS). After incubation with primary antibody, the membranes were washed with chilled 1X PBS thrice and incubated with conjugated secondary antibodies (Donkey anti goat Alexa Fluor^®^ 488, donkey anti Rabbit Alexa Fluor^®^ 594 and DAPI, 1:500) in incubation solution for 1 h in the dark. A no-primary antibody control was included for all experiments. Immunostained samples were imaged on a Nikon C2 confocal microscope (Nikon Instruments Inc., Mellville, NY).

### Western blotting

Protein was isolated from >60 day old hiPSC-RPE cells on transwells using Radioimmunoprecipitation assay (RIPA) lysis buffer (ThermoFisher) along with sonication and was processed as described previously.^1^ The primary antibodies used for Western blotting were anti-Kir7.1 (mouse monoclonal, 1:1000-Santa Cruz Biotech, Dallas, TX), anti-Bestrophin1 (mouse monoclonal, 1:1000 - Novus biologicals, Centennial, CO), anti-RPE65 (mouse monoclonal, 1:1000 - ThermoFisher), anti-GFP (mouse monoclonal, 1:1000-NeuroMab, Davis, CA), both anti-GAPDH (rabbit monoclonal, 1:1000) and anti-β-actin (rabbit monoclonal, 1:1000-Cell Signaling Technology, Danvers, MA) as a loading control. Blots were imaged in Odyssey^®^ Imaging system (Lincoln, NE).

### Phagocytosis Assay

The labelled POS (photoreceptor outer segments) were prepared as previously published and fed to hiPSC-RPE cells growing in transwells that had a transepithelial electrical resistance (TEER) of >150 Ωcm^2^.^16; 22^ The cells were fed POS for either 4 h or 24 h after which any POS that had not been phagocytosed were removed by washing the cells 3 times with DMEM media. The cells were incubated for 24 h or 6 da respectively before imaging. The images were captured and analyzed with NIS-Elements using a Nikon C2 confocal microscope (Nikon Instruments Inc., Mellville, NY).

### hiPSC-RPE Transduction

Lentivirus, custom engineered to be devoid of pathogenic elements, and carrying *KCNJ13* gene fused at N – terminal with green fluorescent protein (GFP) under the control of EF1a promoter, was generated by Cyagen Biosciences (Santa Clara, CA, USA) and used for transduction.^23^ LCA-16 hiPSC-RPE monolayer was infected with pLV-EF1α Kir7.1-GFP at an MOI of 200. The cells were cultured for 4-5 days after infection then used for immunocytochemistry and Western blotting (as described above).

### Electrophysiology

Standard whole cell patch clamp on hiPSC-RPE cells, transduced hiPSC-RPE or hiPSC-RPE treated with read-through drug NB84, with distinct apical processes were performed as described elsewhere.^1^ Data acquisition and the holding potential parameters were controlled using the Axopatch 200-B, Digidata 1550 and Clampex software (Axon instruments). During patch clamping, Ringer’s solution was continuously perfused as an external solution and all drugs were suspended through this solution.

### Statistical analysis

The statistical analysis was performed using Origin (version 9.1) with a two-tailed Student’s *t*-test to assess the significant differences. *P*<0.05 was considered statistically significant. ANOVA and post hoc Tukey test were also used for multiple comparisons. The data are expressed as the means ± *SEM*.

## RESULTS

Analysis of patient derived hiPSC-RPE cells, herein referred to as LCA16 hiPSC-RPE, showed normal RPE morphology, including a cobblestone appearance and pigmentation (Fig. 1a and b), **and was** identical to homozygous wild-type hiPSC **(Fig. S1a**). DNA sequencing confirmed that the control cells were from a heterozygous carrier, herein referred to as control (Ctrl) hiPSC-RPE, while the LCA16 pathogenic variant cells were apparently homozygous for the mutation c.158G>A. The LCA16 hiPSC cells had a normal karyotype (Fig. 1c) confirming that there was no chromosomal abnormality that might affect cellular behavior or response to therapy. The mutation (c.158G>A) introduced a restriction site for Nhe1, enabling the mutant sequence to be identified in patient-specific hiPSC-RPE, further verifying the presence of an apparently homozygous mutation (Fig. 1d). The control hiPSC-RPE cells were analyzed similarly and found to be carrying an undigested 1083 bp band and two digested bands, consistent with the genotype of the donor (Fig. 1d) compared to a single full length transcript band of 1083 bp found in both wild type unrelated hiPSC-RPE cells and commercially available hfRPE cells (Fig. 1d right panel). We believe there was no nonsense-mediated decay in LCA16 hiPSC-RPE as transcript levels were comparable to those in both carrier and wild-type hiPSC-RPE (**Fig. S1c, d**). There was no difference in the expression of RPE-specific genes between the two cell types (Fig. 1e). Thus, the patient-derived iPSC-RPE conformed to the genotype of inherited retinal dystrophy and, therefore, provided a source to establish disease-specific cellular model.^1^

**Fig. 1.**
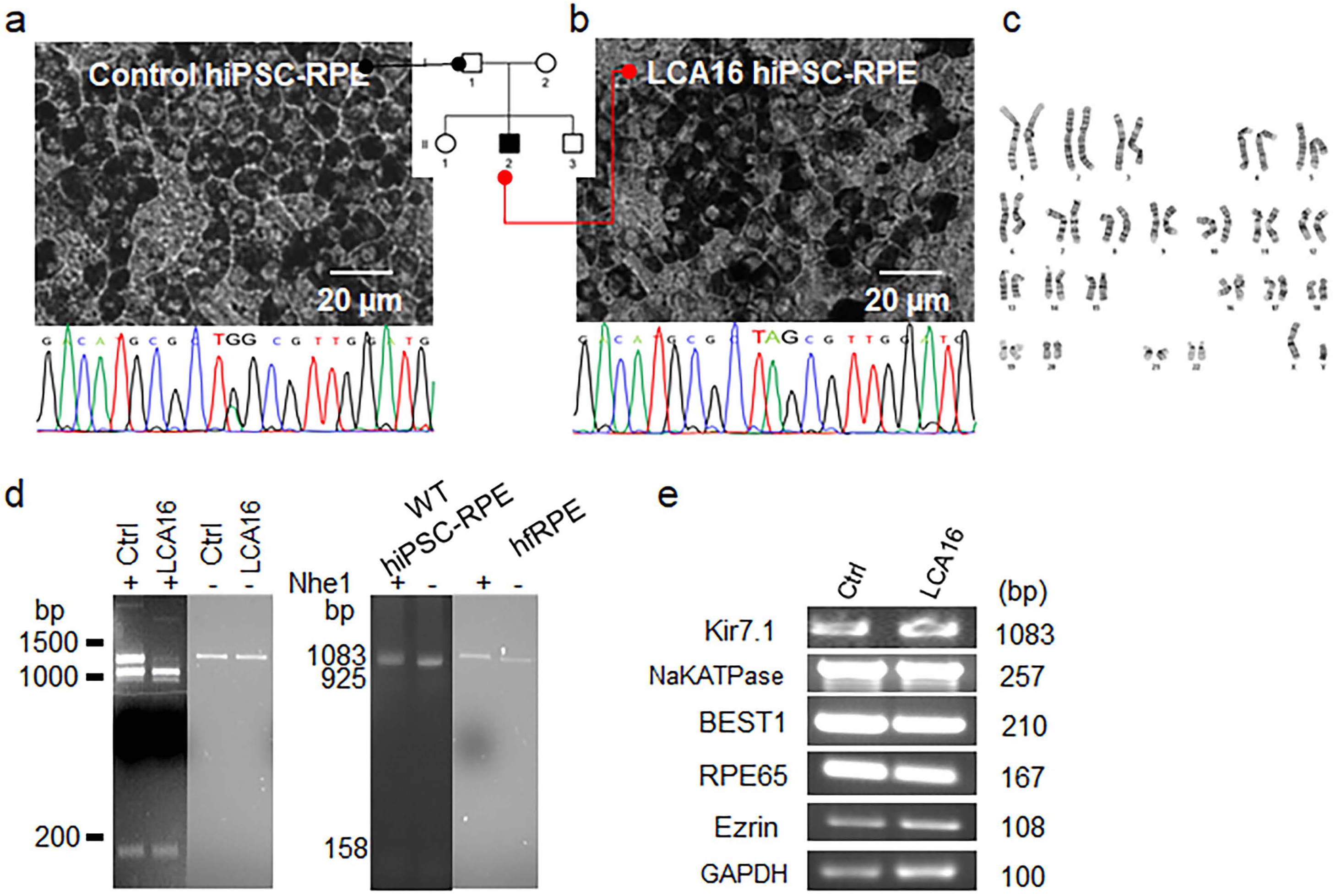
Patient-derived hiPSC-RPE with the LCA16 phenotype. Bright-field images of mature hiPSC-RPE cell monolayer derived from (a) unaffected family member showing heterozygous sequence, and (b) an LCA16 proband with the TAG sequence. Pedigree chart shows sources of hiPSCs. (**c**) Normal karyotype in the patient hiPSC line showing no chromosomal abnormality. (**d**) Analysis of the Nhe1 digestion product from the control, LCA16, and wild-type hiPSC-RPE cells and human fetal RPE cells. The full-length Kir7.1 sequence is 1083 bp in length, and the digested products are 925 and 158 bp in length. (**e**) RPE-specific gene expression in hiPSC-RPE cells.

Kir7.1 channels are located within the highly specialized apical membrane processes of RPE^8; 9^. We therefore examined intactness of apical membrane structures through electron microscope image analysis. The hiPSC-RPE cells had a polarized structure, including intact basal membrane infoldings and elongated apical processes (ap) that measured 1.49 ± 0.05 µm in length in the control hiPSC-RPE and 1.50 ± 0.14 µm in length in the proband’s hiPSC-RPE (P = 0.96, n = 7) (Fig. 2a, c **and Fig. S2**). The distribution and number of mitochondria (m) in the two cell lines appeared normal, averaging 8.4 ± 1.0 and 6.2 ± 0.8 in the control and LCA16 proband’s hiPSC-RPE, respectively (P = 0.12, n = 6) (Fig. 2a and b). Pigment granules correctly distributed apically in both cell types. Kir7.1 protein expression was detected on the apical membrane of mature control hiPSC-RPE cells but not in LCA16 hiPSC-RPE cells (Fig. 2d and e). Western blot analysis for the C-terminal end of the Kir7.1 protein revealed a complete lack of Kir7.1 protein in the LCA16 disease model, as opposed to other RPE proteins that showed equal expression between control hiPSC-RPE and LCA16 hiPSC-RPE (Fig. 2f). The p.Trp53* locus is located within the second exon of the 3-exon *KCNJ13* sequence. We have shown previously that a nonsense substitution at amino acid 53 results in a truncated protein product making the LCA16 patient hiPSC-RPE a null allele, with no Kir7.1 channel function.^1^

**Fig. 2.**
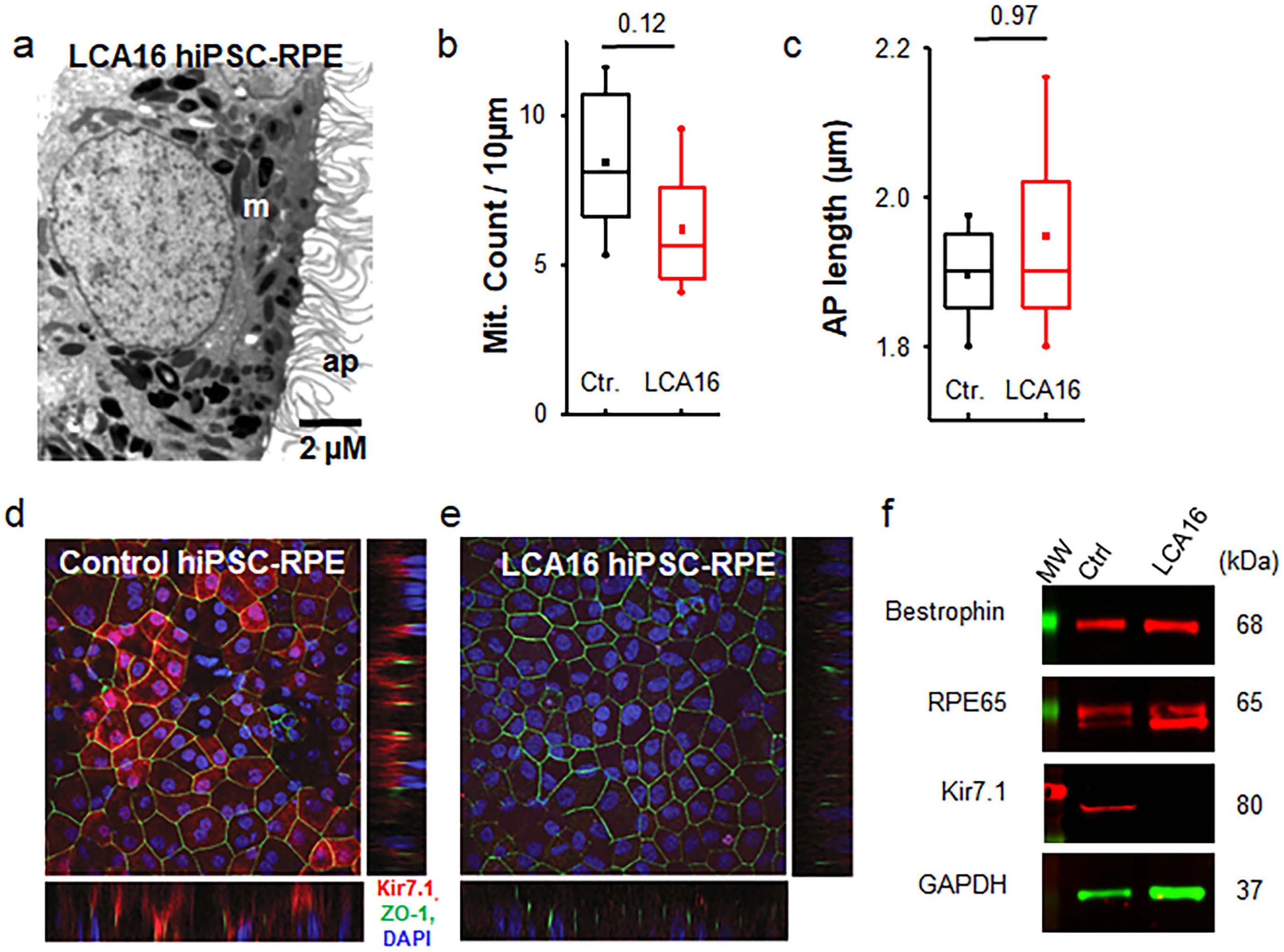
Impaired Kir7.1 protein expression in LCA16 hiPSC-RPE. (**a**) Electron micrograph of a representative LCA16 iPSC-RPE cell. (**b**) Comparison of average mitochondria (Mit.) count within 10 µm^2^ of the cell. (**c**) Evaluation of average length of RPE apical (AP) processes. (**d**) Immunofluorescence localization of Kir7.1 (red), ZO-1 (green) and DAPI (blue) in control hiPSC-RPE cells. Both the lower and side panels reveal a polarized distribution of Kir7.1 in reference to ZO-1 and DAPI (confocal z-stack images). (**e**) Localization of Kir7.1 (red), ZO-1 (green) and DAPI (blue) in LCA16 hiPSC-RPE. (**f**) Western blot showing the expression of RPE cell-specific proteins in both tissue samples. Using a C terminal-specific antibody against Kir7.1, we detected Kir7.1 protein in whole-cell lysates from the control hiPSC-RPE but not in those from the LCA16 iPSC-RPE.

One of the key physiological functions of RPE cells is the daily phagocytosis of the photoreceptor outer segment (POS), which contributes to the POS renewal process. To test whether the absence of normal Kir7.1 protein alters phagocytosis, we fed both control and LCA16 hiPSC-RPE cell cultures with fluorescently labeled photoreceptor outer segments (POS). The cells were fed labeled POS for 4 h, and phagosome digestion by RPE cells was allowed for an additional 24 h. Control hiPSC-RPE showed a higher rate of phagosomal uptake than LCA16 patient hiPSC-RPE (169 ± 40 vs 66.5 ± 7.4, P = 0.04, n = 4) (Fig. 3a, b and c). In addition, when cells were fed with POS for 1 day and then allowed to process phagosomes for 6 days, the LCA16 hiPSC-RPE cells failed to digest POS (80.2 ± 11.1 vs 244.2 ± 27.6 counts within a 200 µm^2^ field, P = 0.001, n = 4) (**Fig. S2**). This finding suggests that the fundus pigmentation observed in LCA16 patients is likely due to an inability to phagocytose POS normally, which therefore accumulate over time in the retinas of affected individuals.

**Fig. 3.**
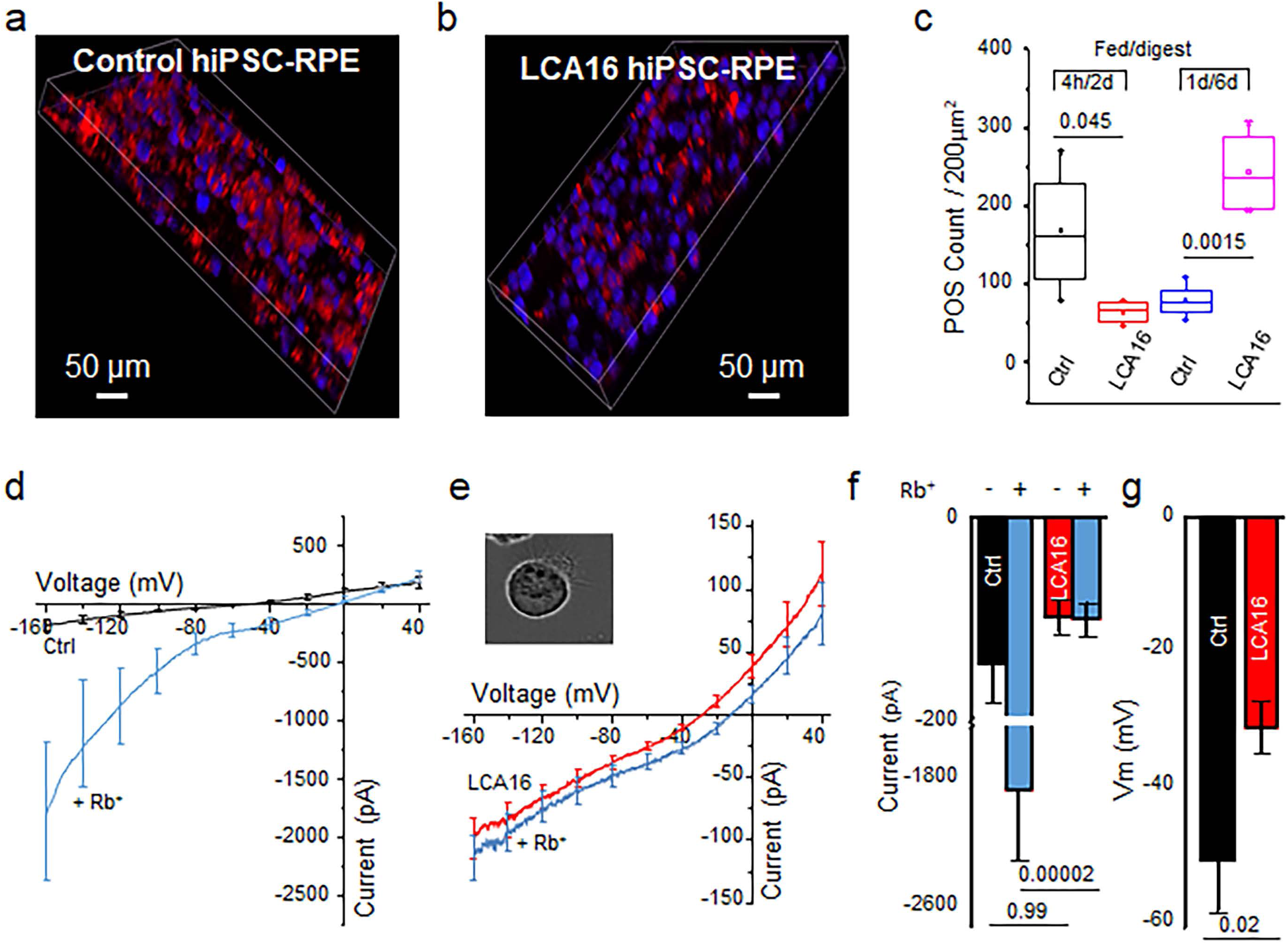
LCA16 hiPSC-RPE exhibit aberrant physiology. Phagosome (red) localization within control hiPSC-RPE (**a**) and LCA16 iPSC-RPE (**b**) samples. (**c**) Plot of the average phagosome count within a fixed 200 µm^2^ area in the control and LCA16 hiPSC-RPE cells after 4 hrs of feeding and a subsequent 48-hr digestion period, or after 1 day of feeding followed by 6 days of digestion. (**d**) Plot of the average current-voltage (I/V) curve for Kir7.1 currents using normal external K+ (black) or high external Rb+ (light blue) in control hiPSC-RPE cells. (**e**) An average I/V curve using normal K+ (red) and high Rb+ (light blue) in LCA16 hiPSC-RPE cells. (**f**) Average plot of an inward current amplitude measured at −150 mV. Color representation as shown in (d) and (e). (**g**) Comparison of the average membrane potential of the control (black) cells to depolarized LCA16 (red) hiPSC-RPE cells.

By using differentiated hiPSC-RPE from an asymptomatic carrier family member that develop apical processes and localize Kir7.1 to the apical membrane, we were able to detect a small but measurable Kir7.1 current (−144.2 ± 51 pA) in control hiPSC-RPE cells. Normal function was confirmed by a 13-fold increase in Rb+ permeability (−1887.3 ± 589.6 pA, n = 5) (Fig. 3d and f, **similar to wild-type hiPSC Fig. S1b**), which is a specific property of the Kir7.1 channel.^1; 24^ However, in LCA16 hiPSC-RPE cells, we did not detect any change in the current amplitude mediated by Rb+ conductance (−98.1 ± 15.7 pA & −99.6 ± 15.7 pA, n = 9) (Fig. 3e and f). A direct comparison of both current amplitude (P = 0.0006 with Rb+) and cell membrane potential (−50.0 ± 5.1 vs −30.6 ± 3.7 mV, P = 0.0005; as shown in Fig. 3g) supports our hypothesis that the cause of blindness is a nonfunctional Kir7.1 channel which depolarizes LCA16 hiPSC-RPE cells. This is similar to observations in an *in vivo* Kir7.1-deficient mouse model or an *in vitro* cell line exogenously expressing the human Trp53* Kir7.1 channel.^1; 9^

We next assessed the functional consequences of the NB84-mediated read-through on Kir7.1 current in LCA16 hiPSC-RPE cells. Following treatment of the LCA16 hiPSC-RPE with 500 µM NB84, we obtained a measurable current of −94.3 ± 24.0 pA, which was enhanced 10-fold upon the introduction of the permeant ion Rb+ (−1562.7 ± 546.7 pA, P = 0.005, n = 8; Fig. 4a and b). A significant recovery of membrane potential from −30.6 ± 3.7 in the untreated cells to –56.3 ± 3.6 mV (P = 0.0001, n = 10; Fig. 4c) in the NB84 treated cells further supports the potential of read-through drug therapy for restoring channel function. In a subgroup of cells, we noticed rescue in membrane potential without any significant change in current amplitude. One possible explanation for this finding is the heterogeneity in the incorporation of the near-cognate amino acid during Kir7.1 protein translation (**Fig. S3**).

**Fig. 4.**
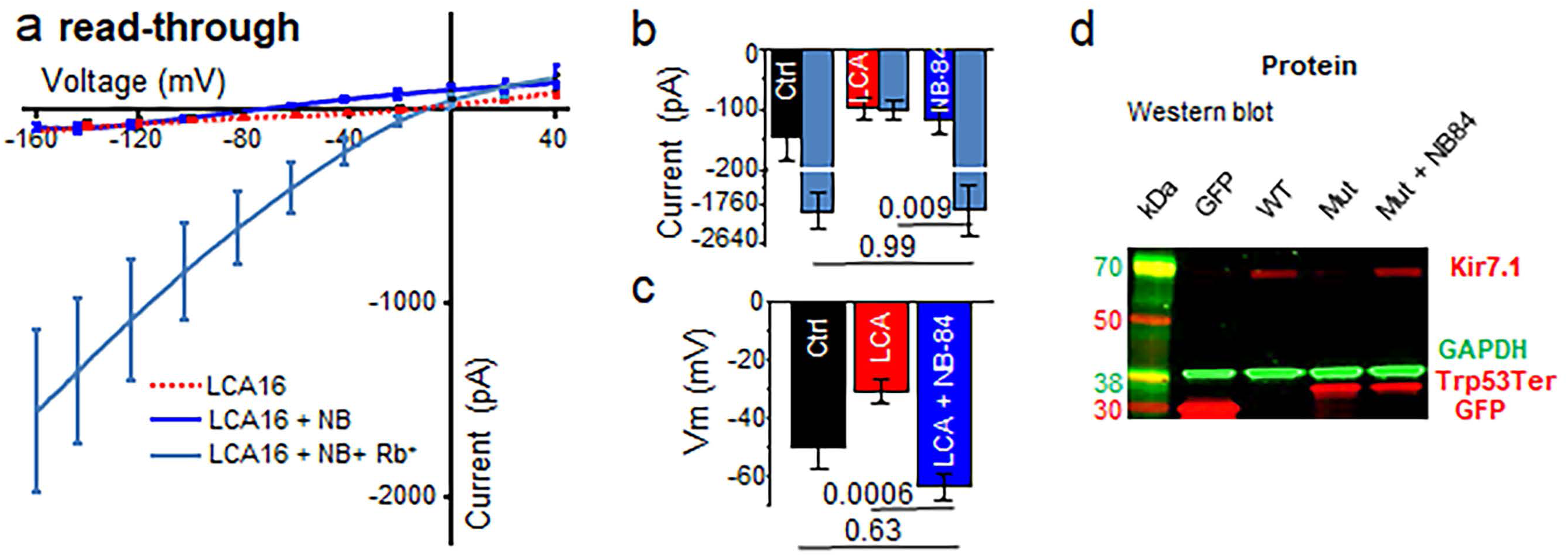
Putative Kir7.1 loss-of-function rescue through nonsense suppression. (**a**) Average I/V relationship before (red) and after (dark blue) treatment with NB84. The Rb+ current measured is shown as a light-blue trace. Evaluation of the average inward current measured at −150 mV (**b**) and of the membrane potential (**c**) to demonstrate the effect of NB84. (**d**) Western blot showing detection of eGFP fusion proteins, using anti-GFP antibody, in cell lysates from CHO cells transfected with empty vector (GFP), WT Kir7.1 coding sequence (WT) or Trp53* coding sequence (Mut). A partial restoration of the full-length protein product was observed in the latter after NB84 treatment (Mut + NB84).

Endogenous Kir7.1 protein levels in the NB84 treated hiPSC-RPE were below the detection level of Western blots so we transfected CHO cells with a mutant Kir7.1-eGFP fusion construct to detect the formation of the full length protein. After treating these cells with NB84, full length protein in addition to truncated protein was detected on a Western blot and both membrane potential and current deficits were corrected (Fig. 4d and **Fig. S4**), demonstrating that NB84 potentiates the specific read-through of the recessive Trp53* mutation.

We designed a lentiviral vector with an N-terminal GFP fused to the wild-type human Kir7.1 open reading frame under the control of the EF1a promoter,^25^ which was subsequently used to infect hiPSC-RPE monolayers. Intriguingly, wild-type Kir7.1-expressing cells presented normal Kir7.1 currents or even slightly higher amplitudes than those observed in the control cells (−921 ± 223 pA, P = 0.001, n = 8). This current was further potentiated by the introduction of Rb+ (−5453 ± 929 pA), as expected for a normally functioning Kir7.1 channel (Fig. 5a and b). In addition to K+ currents, the membrane potential of LCA16 iPSC-RPE cells was approaching normal values (−57.5 ± 5.4 mV, P = 0.0008) (Fig. 5c). Moreover, newly expressed full length Kir7.1 (Fig. 5d) was shown to be localized to the apical membranes of patient-specific hiPSC-RPE cells (Fig. 5e **and Supple. Video**).

**Fig. 5.**
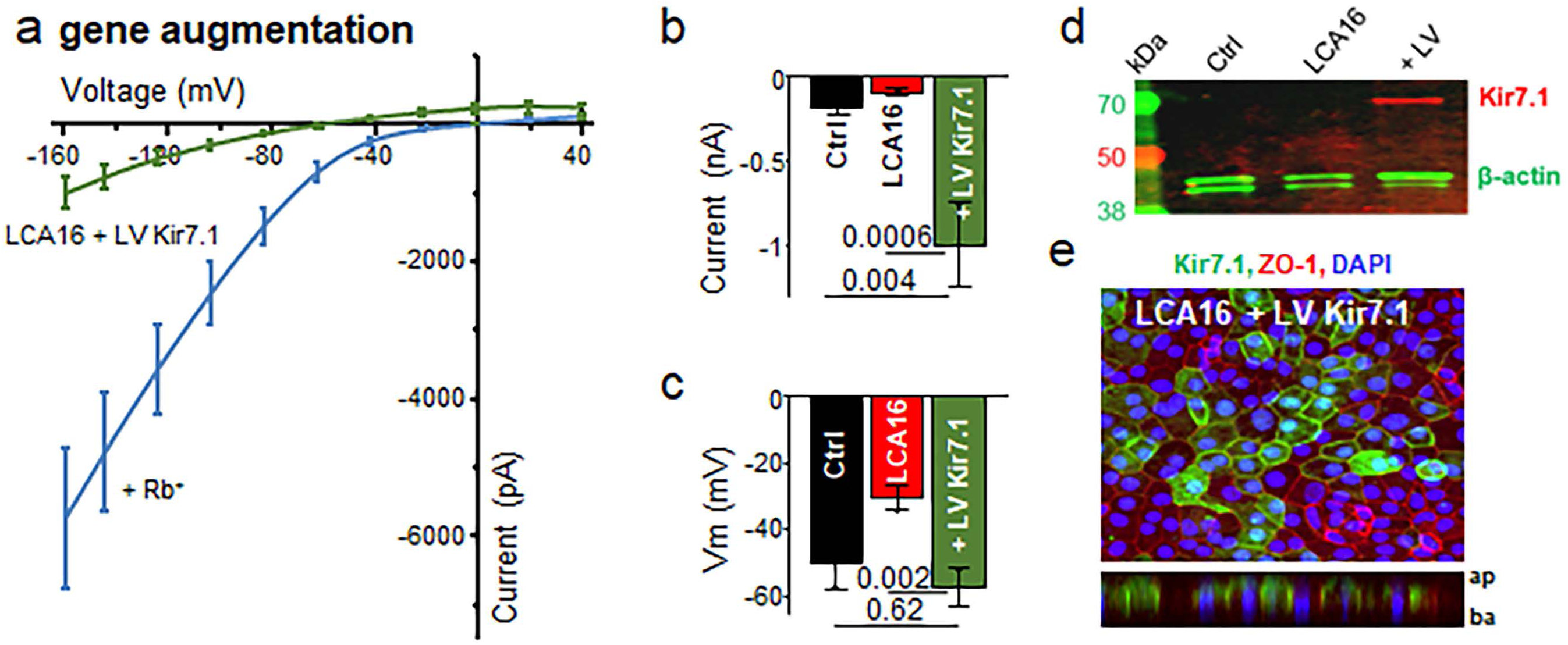
hiPSC-RPE regain Kir7.1 function following gene-augmentation. (**a**) Plot of the average I/V curve for Kir7.1 currents measured in LCA16 hiPSC-RPE GFP-positive cells expressing a normal copy of the human Kir7.1 clone. Both K+ (green) and Rb+ (light blue) traces are shown. Average plot of the (**b**) current amplitudes measured at −150 mV and (**c**) membrane potential show rescue after gene augmentation. (**d**) Western blot analysis of lysate from control hiPSC-RPE (Ctrl), LCA16 hiPSC-RPE (LCA16) or LCA16 hiPSC-RPE infected with lentivirus coding for the eGFP-WT Kir7.1 fusion protein (LV) using anti-GFP antibody. (**e**) Cultured LCA16 hiPSC-RPE showing wild-type Kir7.1 (green), ZO-1 (red) and DAPI (blue) proteins. Z-stack planes shown in the lowerpanel.

## DISCUSSION

We have reported previously that targeted inhibition of Kir7.1 in the mouse retina (induced using either siRNA or a pharmacological blocker) causes an altered electroretinogram phenotype consistent with that observed in LCA16 patients.^9^ In the current report, we extend our findings by describing the development of a novel LCA16 disease-in-a-dish model. We developed a robust hiPSC-RPE cell culture derived from skin biopsies from one LCA16 proband carrying an apparently homozygous nonsense mutation in exon 2 of the *KCNJ13* gene (p.Trp53*) and a control from an unaffected healthy family member.^16^ The highly pigmented RPE cells from both control and patient appeared normal, expressed RPE markers and were immunopositive for several RPE specific proteins except the LCA16 hiPSC-RPE lacked mutation-specific Kir7.1 protein expression. Rescue of Kir7.1 function in RPE cells shown by us is an intervention that could improve vision in patients with congenital blindness due to *KCNJ13* mutations.

Our study determined that the major pathogenic alteration affecting the RPE cells is the lack of functional Kir7.1 channel. We were also able to establish a genotype-phenotype (normal) correlation in an asymptomatic carrier individual suggesting complete penetrance of p.Trp53* pathological mutation. Use of carrier (control) hiPSC-RPE, from within the family, also accounts for action of modifier genes, epigenetic changes or environmental contribution. We have previously determined that cultured human fetal RPE cells express Rb+ sensitive Kir7.1 current similar to what we recorded from wild-type and carrier hiPSC-RPE cells.^19^ In our hands, heterozygous expression of equal amounts of wild type and non-functional p.Trp53* mutant protein also does not influence cellular physiology outcome.^1^ Given these findings, we reasoned that carrier hiPSC-RPE are a suitable control to compare test results of both gene restoration and translational read-through.

Our diseased hiPSC-RPE model is particularly valuable given that a *Kcnj13* gene knock out in mice has been proven to cause embryonic lethality suggesting that there is a different function of *Kcnj13* gene in mice or animal models that complicates the study of disease pathobiology.^12; 26; 27^ We showed that patient-specific hiPSC-RPE cells had normal apical processes where Kir7.1 protein is expressed. We thus confirmed maturity of these cultured hiPSC-RPE cells unlike the ones generated from ciliopathy patients.^28^ Matured LCA16 hiPSC-RPE cells did not show Kir7.1 function, their membrane potential was depolarized and they were unable to phagocytose photoreceptor outer segments as compared to control hiPSC-RPE cells. A defect in the phagocytosis of shed photoreceptor outer segments by LCA16 hiPSC-RPE is thus secondary and consistent with the slow progression toward blindness observed in LCA16 patients, as documented through clinical manifestations, including electroretinogram abnormalities and retinal pigmentation.^1–4^ In general, it is challenging to study ion-channels in iPSCs due to the lack or low level of expression but we have used the powerful tool of stem cell technology to analyze the pathophysiology of LCA16.^29^

The LCA16 mutation p.Trp53* analyzed in this report is a tryptophan (UGG) to amber (UAG) stop codon variant.^1^ An in-frame nonsense mutation accounts for about 12% of inherited genetic disorder.^30^ Several studies have demonstrated the usefulness of read-through therapy as an intervention that could improve vision.^31–33^ Designer aminoglycosides such as NB84 bind to eukaryotic ribosomal RNA decoding site.^34^ This binding promotes the transition of the 16S rRNA decoding center from a tRNA binding conformation to a productive state resulting in an increase in the rate of read through at a stop codon.^35^. Here we show that a *KCNJ13* p.Trp53* nonsense mutation is suppressed by the incorporation of a near-cognate aminoacyl tRNA in the presence of the small-molecule read-through designer aminoglycoside NB84.^33; 36; 37^ Aminoglycosides pose toxicity concerns in the adult, but both *in vitro* and *in vivo* toxicity studies demonstrate that NB84 is ready for translational therapy in animal models and human subjects.^38^ We foresee further development of a non-toxic formulation of NB84 (Poly-L-aspartate that increases cytoplasmic concentration^39^) for topical application to eye so that risks to other organs are avoided. As expected, only a fraction of cells showed complete recovery of both membrane potential and current amplitude whereas there was a subgroup with partial recovery of function. Read-through therapy also has a significant clinical advantage in the management of LCA16 at an early stage when diagnosed.

The particular mutation studied herein, and other recessive mutations that cause blindness, are potential targets for gene therapy. This premise is nearing reality given the recent FDA approval of a gene therapy treatment for blindness.^40,41; 42^ Using a LCA16 hiPSC-RPE model, we demonstrate that the rescue of channelopathy via lentiviral gene augmentation produced a potassium current and normalized membrane potential (**Fig. S5**). Newly expressed protein was also localized to apical aspects of LCA16 hiPSC RPE cells.

Gene therapy is a very simple concept but the problem remains that too little or too much of the expression will have deleterious effects.^43; 44^ To understand an optimum dosing required in our case, we expressed copies of both wildtype and mutant protein in a heterologous expression system. We observed that as little as 25% of wildtype protein expression of total Kir7.1 protein product was sufficient to enhance both membrane potential as well as potassium current and rescued the mutant phenotype (**Fig. S6**). A splice variant of Kir7.1 was reported in human RPE cells which failed to produce anticipated small protein band.^45^ Furthermore, the expected truncated protein consisted of N-terminus and the first transmembrane domain so the LCA16 nonsense mutation p.TRP53W* is predicted to truncate all Kir7.1 transcript variants. The p.Trp53* truncated protein product does not assemble with wild type protein to alter Kir7.1 channel function and does not perhaps undergo nonsense-mediated decay as a truncated protein product is expressed after heterologous expression.^1^

In this study we have taken advantage of long lasting transgene expression in RPE cells *in vivo* through Lentivirus transduction.^46^ Also, AAV in our hand was unable to transduce iPSC-RPE cells as has been experienced by several other groups. This might be due to the higher efficacy of lentiviral vector to transduce post mitotic RPE cells as has been proved for non-dividing cells.^47^ Our 3^rd^-generation lentivirus packaging system also has reduced risk of generating replication-competent virus. In a preclinical study carried out in non-human primate, subretinal delivery of lentiviral RPE65 gene showed that the vector stayed in the eye perhaps because of the tight blood-retina barrier with high systemic safety.^48^ Retinal gene therapy using lentiviral vector is in clinical trials for Neovascular age-related macular degeneration, Stargardt disease, and Usher syndrome type 1B.^49^ We conclude that both treatments developed in this study are able to achieve at least 25% normal protein expression as restoration of membrane potential was complete with variable recovery in current amplitude. Achieving this 25% functional rescue *in vivo* will thus be less challenging and mutant protein product will not influence functional outcome.

Advances in genetic screening and improvements in access to medical care will undoubtedly improve our understanding of the array of disorders caused by channelopathies and expand our understanding of the role that *KCNJ13* plays in health and disease. Genome editing holds promise for genetic disorders but gene correction in non-dividing cells and introduction of unwanted genomic variants at the target and off-target sites limits its potential use. Overall, we show a promising proof of concept for read-through and gene-augmentation therapy using a precision medicine approach that would benefit pediatric blindness (LCA16) patients.

## Supporting information

## Acknowledgments

We sincerely thank the patients and their family members for their willingness to participate and donate biopsy skin samples and time. We also thank the anonymous gift for LCA16 research through the UW-Foundation. NB84 was kindly provided by Ellen Welch. This work was also supported by an NEI grant (EY024995-BRP), UW Vision Core Grant P30 EY01665, and the Retina Research Foundation M. D. Mathews Professorship (BRP) at the UW-School of Medicine and Public Health and Department of Pediatrics. DMG is supported by the McPherson Eye Research Institute Sandra Lemke Trout Chair in Eye Research and the Retina Research Foundation Emmett Humble Distinguished Directorship. We thank Kris Saha for his suggestions on this manuscript and Randall J. Massey for help with electron microscopy. An unrestricted grant from Research to prevent Blindness to the UW Madison Department of Ophthalmology and Visual Sciences also supported this work. The views of this report are primarily of the authors and were not influenced by the funding agencies. The members of the Pattnaik Lab are also recognized for their input. Present address of DMP is Department of Pediatrics, University of Illinois Chicago, Chicago, IL.

## Author contributions

BRP initiated and supervised this project. BRP, DMG and DMP conceived the study. DS, KDB, EC and DMG helped generate hiPSC-RPE lines. BRP and PKS generated and analyzed electrophysiology data. PKS, DH, SB, HM and SS generated and analyzed the molecular biology data. BRP and PKS generated the figures and wrote the manuscript. All of the authors have read, edited and approved the manuscript.

