## Supplementary material for "Gene augmentation and read-through rescue channelopathy in an iPSC-RPE model of congenital blindness"

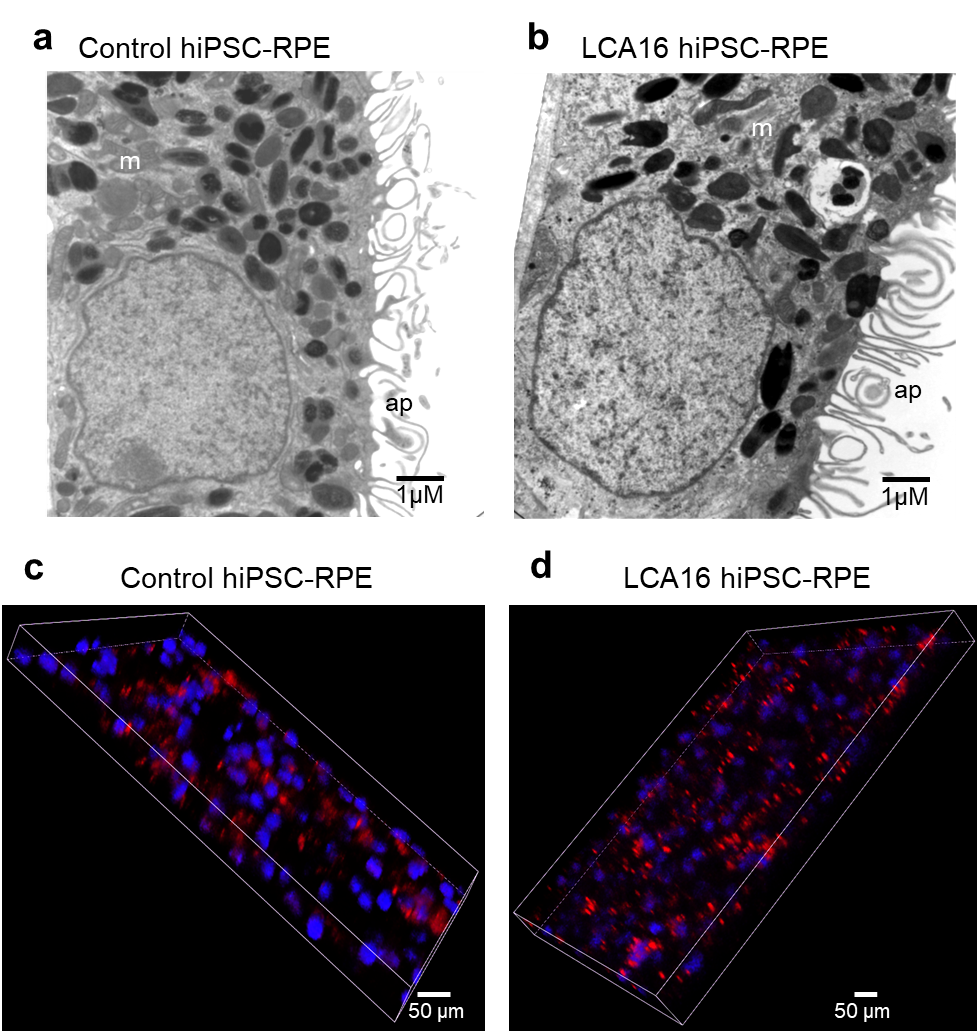

**Supplemental Figure 1**

**Phenotype of patient-derived iPSC-RPE cells.**

Comparison of electron micrograph of a control hiPSC-RPE cell (a) and an LCA16 hiPSC-RPE cell (b) showing normal columnar morphology with basal infoldings, large nuclei, mitochondria (m), melanosomes and intact apical membrane with extending processes (ap). Images of live control hiPSC-RPE (c) and patient-derived hiPSC-RPE (d) cells in x-y-z dimension showing POS (red) and nuclei (blue) imaged 6 days after feeding cells for 1 day with fluorescent-labeled bovine POS. More undigested red fluorescent POS particles are visible in LCA16 hiPSC-RPE cells.

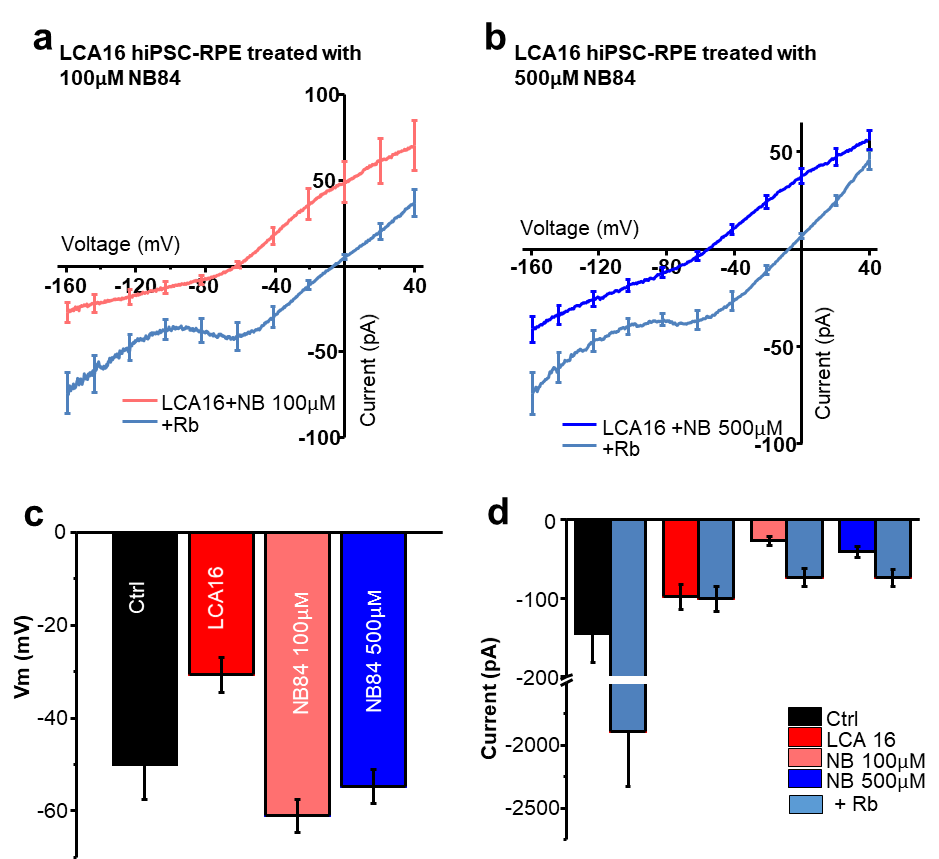

**Supplemental Figure 2**

**Subpopulation of hiPSC-RPE show rescue in membrane potential but not current amplitude.**

a) I/V plot of average current response in a subgroup of LCA16 hiPSC-RPE cells in normal K+ Ringer’s and high Rb+ Ringer’s solution after treatment of cells with 100 µM NB84. b) I/V plot showing K+ and Rb+ current response in LCA16 hiPSC-RPE after treatment with 500 µM NB84. c) Average plot of membrane potential showing rescue of membrane potential to control levels after treatment of LCA16 hiPSC-RPE with 100 or 500 µM NB84. d) Current amplitude plot clearly demonstrating no rescue in current amplitude after treatment with either 100 or 500 µM NB84.

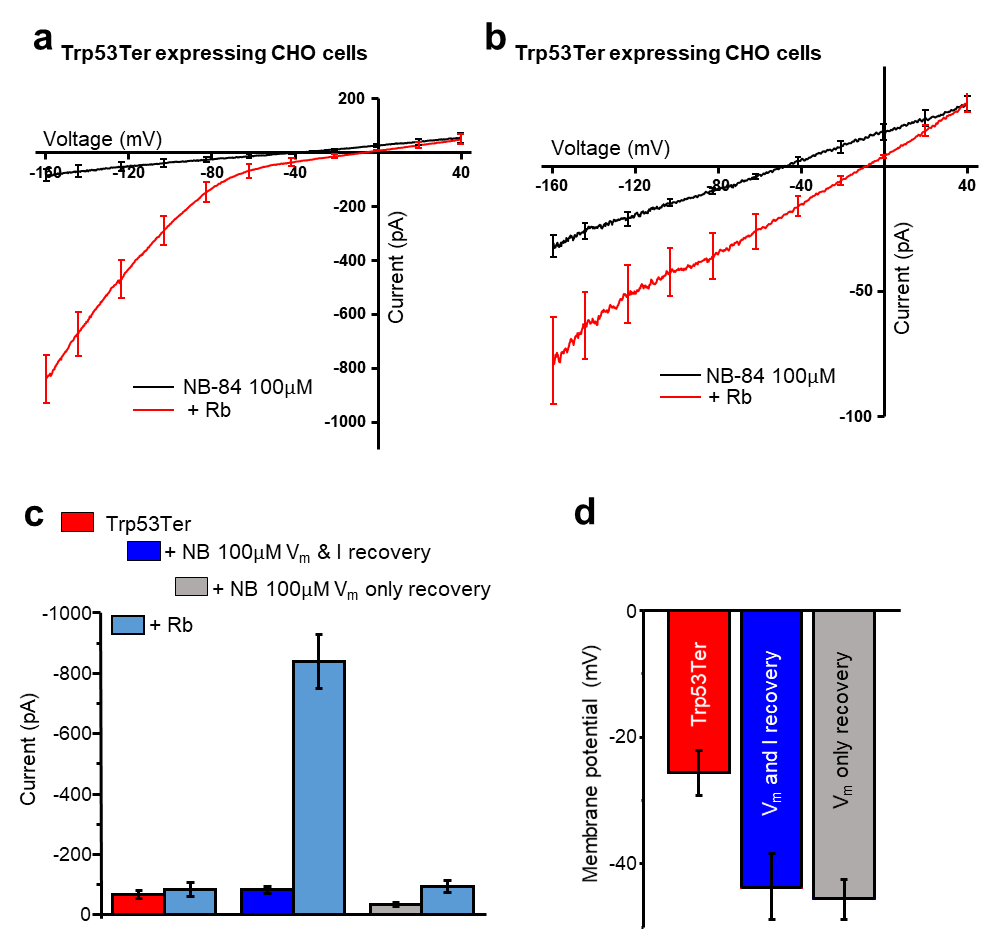

**Supplemental Figure 3**

**Read-through of Trp53Ter ectopically expressed in CHO cells.** As in LCA16 hiPSC-RPE cells, transduced CHO cells showed inwardly rectifying Kir7.1 current activated by Rb+ only after treatment with NB84. a) I/V plot of cells showing K+ (black) and Rb+ (red) current after treatment with NB84 showing recovery of both current amplitude and membrane potential. b) A group of NB84 treated cells showing somewhat linear I/V plot for K+ (black) and Rb+ (red) illustrating recovery of only membrane potential but not current amplitude. Comparison of average recovery of both current amplitude (c) and membrane potential (d) after treatment of Trp53Ter expressing CHO cells.

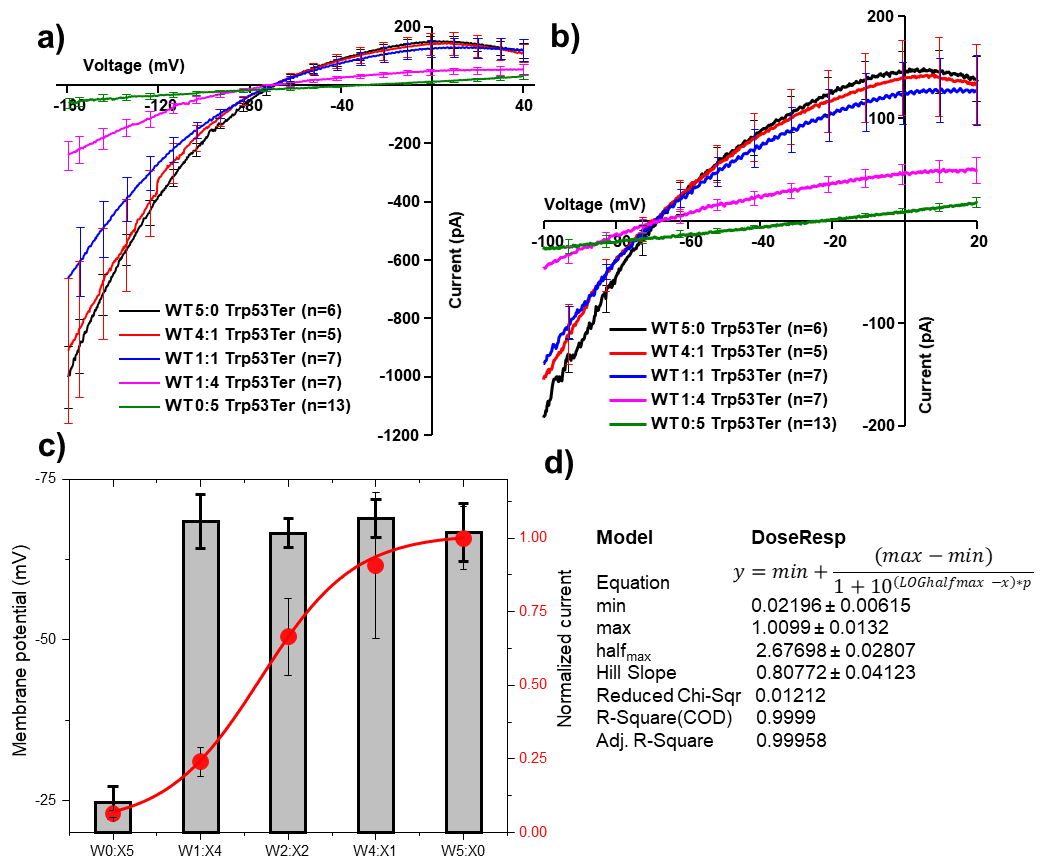

**Supplemental Figure 5**

**Determination of the extent of wildtype protein expression required for functional rescue.** We were particularly interested in quantitating how much gene augmentation/correction is required to restore channel function. We expressed either Trp53Ter or wild type Kir7.1 protein alone or in various combinations in CHO cells. a) Current recordings are shown as I/V plots. b) On an expanded scale for x-axis, resting membrane potential shows negative shift with wild type protein making up only 20% of the protein expression. c) Average plot of either normalized current amplitude (filled circles) or membrane potential (grey bar) as a function of increasing wildtype protein expression. Solid line is a best fit for distribution using equation shown on the right. Half-maximum current was obtained with about 26% of the wild type protein expression.

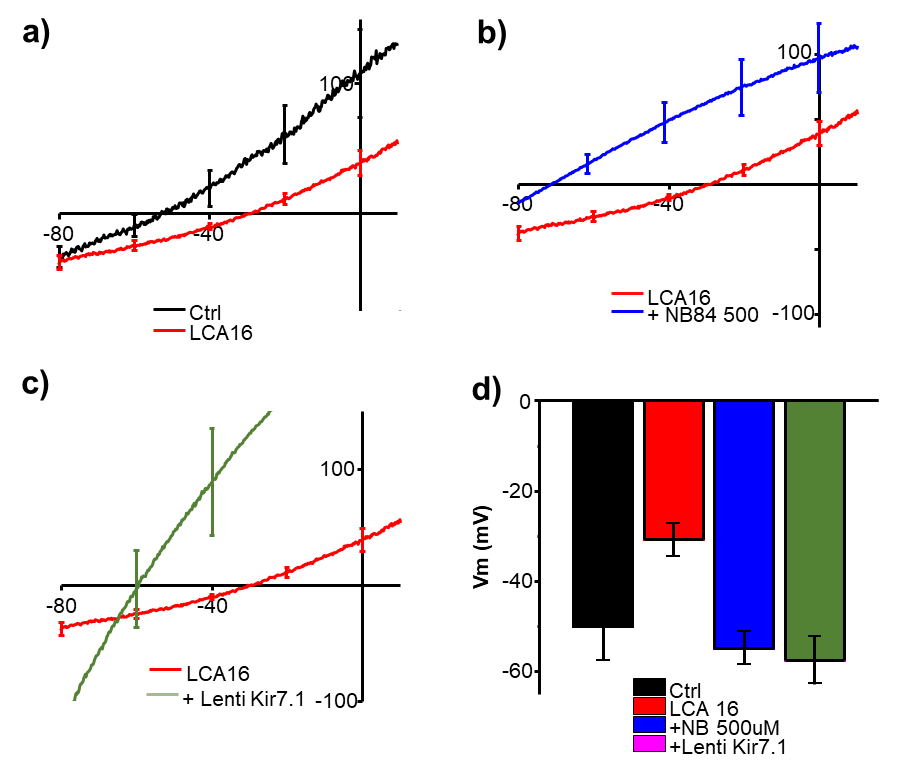

**Supplemental Figure 4**

**Comparison of the Rescue of membrane potential across treatment modalities.** a) On an expanded scale of the x-axis, resting membrane potential of control (black) and LCA16 iPSC-RPE (red) showed a positive shift in I/V plot. b) For the LCA16 iPSC-RPE cells (red trace), treatment with NB84 shifted the I/V-plot to negative (blue). c) Plot of average I/V also showed a negative shift of resting potential after gene augmentation (green). d) Bar graph comparison of resting membrane potential showed recovery of LCA16 iPSC-RPE to control level after treatment with NB84 or upon gene augmentation.
